# Integrative identification of *Tetratostichococcus* sp. CCAH0201 (Trebouxiophyceae, Chlorophyta) from *Rhizophora* leaves in the Can Gio Mangrove Forest, Vietnam

**DOI:** 10.64898/2026.11.05.764477

**Authors:** Thanh-Luan Nguyen, Hoang-Dung Tran

## Abstract

A microalgal strain from *Rhizophora* leaves in the Can Gio Mangrove Forest, Vietnam, was examined by light microscopy and three molecular markers. Thirty-six cells from 22 micrographs measured 6.68–12.74 × 3.49–4.79 µm. Homologous phylogenetic windows comprised 550 nucleotides for SSU rDNA, 711 for ITS and 1,170 for *rbc*L. CCAH0201, P1 and *Tetratostichococcus jenerensis* SAG 2138 shared the SSU-3′ sequence, whereas ITS and *rbc*L placed CCAH0201 with P1 (SH-aLRT/UFBoot 87/94 and 96.9/99). CCAH0201 differed from P1 at 22/711 ITS and 4/1,170 *rbc*L sites. Structure-guided ITS2 comparison identified one strict CBC, one two-sided change from a wobble pair and three hemi-CBCs relative to P1. CCAH0201 is thus identified as *Tetratostichococcus* sp. in a lineage close to P1.

## 1. Introduction

Leaves from the Can Gio Mangrove Forest, Vietnam, host microalgae associated with plant surfaces. Several *Rhizophora*-leaf isolates in the CCAH collection were initially recorded as *Stichococcus*-like based on cell shape. Because the small cylindrical cells have few distinctive features visible by light microscopy, identification relies on comparisons with authenticated material and homologous sequences. A study of CCAH Desmodesmus strains from Can Gio likewise combined morphology with rDNA markers to refine microalgal identification (Luu et al., 2024).

The revision by Pröschold and Darienko (2020) showed that algae collectively described as *Stichococcus*-like belong to several phylogenetic lineages. In that framework, *Tetratostichococcus* was distinguished using phylogenetic evidence and features of the SSU rDNA V9 region; *T. jenerensis* is the type species of the genus. Cell morphology, cellular organization, and ITS2 characters were considered together rather than applying a sequence-similarity percentage to assign species names. That study also showed that culture conditions can substantially affect morphology in members of this group, limiting the diagnostic value of cell length or length-to-width ratio alone (Pröschold and Darienko, 2020).

The comparative set for *Tetratostichococcus* includes named material and strains still reported at genus level. Strain P1, isolated from tropical peat soil in Malaysia, has genomic data that allow several markers from one material to be compared (Sahabudin et al., 2022; Maeda et al., 2024). A Fergana Valley soil isolate was also retained at genus level after morphological and molecular study (Tukhtaboeva et al., 2024). These strains broaden comparisons with CCAH0201 alongside the type-species material *T. jenerensis* SAG 2138.

ITS2 provides discriminating characters when SSU varies little. Interpretation of compensatory base changes (CBCs) requires homologous paired positions in a structural model. CBC counts depend on how the structure is inferred and how confidently positions are mapped (Caisová et al., 2013; Wolf et al., 2013), especially when some reference material has only short ITS2 fragments.

This study asks whether a *Stichococcus*-like strain from *Rhizophora* leaves can be distinguished from known *Tetratostichococcus* material when SSU remains highly conserved. We compared CCAH0201 with authenticated *T. jenerensis* SAG 2138 (SAG, 2026) and the genome-characterized strain P1 using morphology, nuclear and plastid loci, and ITS2 structure. This approach clarifies the placement of the Can Gio strain and identifies evidence needed for future genome-scale tests.

## 2. Materials and methods

### 2.1. Origin and maintenance of the microalgal strain

*Tetratostichococcus* sp. CCAH0201 was isolated from the surface of *Rhizophora* leaves in the Can Gio Mangrove Forest, Ho Chi Minh City, Vietnam (10.400820° N, 106.904870° E; Fig. 1) and is maintained in the Ho Chi Minh City Collection of Algae (CCAH). The strain is maintained in Bold’s Basal Medium (BBM).

**Figure 1.**
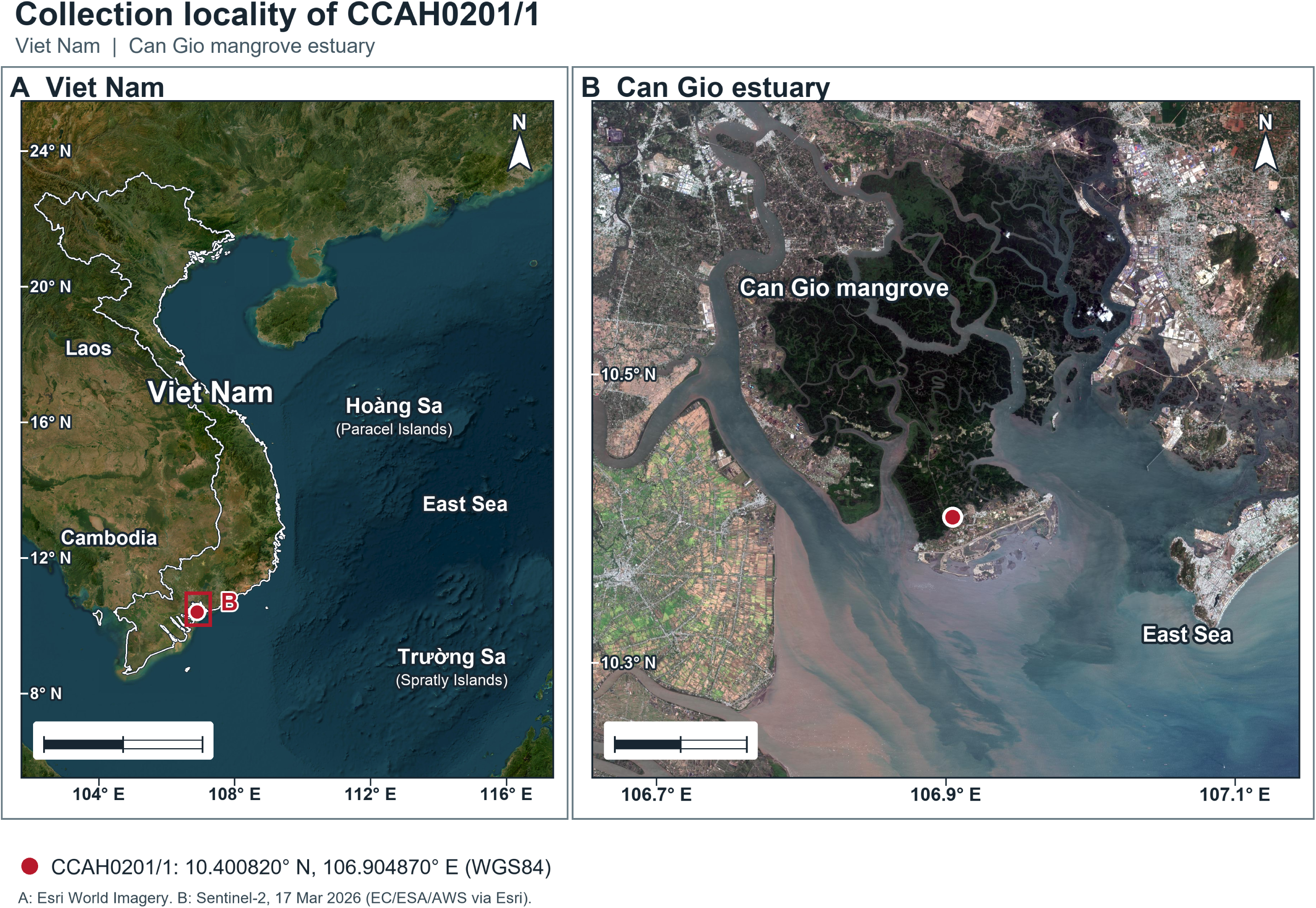
Collection locality of CCAH0201/1 in Can Gio, Ho Chi Minh City, Viet Nam. (A) Viet Nam locator map showing Hoàng Sa (Paracel Islands) and Trường Sa (Spratly Islands); scale bar = 500 km. (B) Natural-colour Sentinel-2 image of the Can Gio estuary acquired on 17 March 2026; scale bar = 10 km. The red dot marks the coordinates in the collection record (10.400820° N, 106.904870° E; WGS84). Panel A imagery: Esri, Vantor, Earthstar Geographics, and the GIS User Community; panel B imagery: Esri, European Commission, European Space Agency, and Amazon Web Services; accessed 4 November 2026. The national outline in panel A is from Natural Earth 1:10m.

### 2.2. Morphological observation and quantification of cell dimensions

Cell morphology of CCAH0201 was examined by light microscopy using 24 archived micrographs. The published CCAH morphological workflow assessed cell shape, size, and arrangement by light microscopy at 40× and 100× (Luu et al., 2024). All 24 CCAH0201 micrographs used for quantification in the present study were labelled 100× in their source files; two image groups labelled 1× and 2× were calibrated separately against their scale bars. The micrographs used for measurement carried a 10-µm scale bar on the image. Scale bars in the two groups measured 552 and 1,083 pixels, corresponding to conversion factors of 0.018116 and 0.009234 µm/pixel, respectively. Figure panels were cropped and arranged from the original micrographs without altering cellular features.

Cells selected for measurement were isolated, intact, had clearly defined boundaries, and did not touch or overlap other cells. Outlines were determined on full-resolution images. The longer and shorter sides of the minimum-area bounding rectangle, rotated to the cell axis, were used as cell length and width, respectively. Objects at image edges or with uncertain outer boundaries were excluded.

The length-to-width ratio was calculated for each cell. Within-observer repeatability was assessed by blinded remeasurement of 10 of 36 cells (27.8%) from coded image crops. Measurements were summarized as mean ± sample standard deviation, median, Q1–Q3, and minimum–maximum range. The dataset comprised 36 cells from 22 micrographs of one strain; cells and micrographs were not treated as independent biological replicates.

### 2.3. DNA extraction, PCR amplification, sequencing, and marker curation

Total DNA was extracted from cultured CCAH0201 biomass and used as the template for PCR. The SSU rDNA and ITS1–5.8S–ITS2–LSU regions were amplified with primers EAF3/ITS055R; internal primers G500F and G800R were used to improve coverage and check both sequencing directions following a workflow applied to *Stichococcus*-like algae (Pröschold and Darienko, 2020).

The plastid *rbcL* region was amplified using the standard primer system PRASF1 (5′- ATGGTTCCACAAACAGAAAC-3′) and PRASR1 (5′- TTGTCAATAGTATCAAATTC-3′); internal primers PRASF2/PRASR2 were used for overlapping reads when needed (Sherwood et al., 2000; Neustupa et al., 2013). Thermal cycling consisted of 95°C for 5 min; 40 cycles of 95°C for 45 s, 50°C for 90 s, and 72°C for 2 min; and a final extension at 72°C for 10 min.

PCR products were checked by electrophoresis in 1% agarose, purified and sent to a sequencing service for Sanger sequencing. Returned sequences were checked for strain and marker identity, read orientation and ambiguous characters before inclusion in the analysis matrices. The *rbc*L fragment was also checked for reading frame and internal stop codons.

The CCAH0201 sequence set comprised a 600-nucleotide SSU segment, an 809- nucleotide ITS sequence, and a 1,425-nucleotide *rbc*L fragment. The *rbc*L fragment contained 475 codons and no internal stop codon in the checked reading frame. Homologous windows used for phylogenetic analysis were 550 nucleotides for SSU-3′, 711 for ITS, and 1,170 for *rbc*L.

### 2.4. Sequence standardization and analysis datasets

GenBank assigned QB266712 to the 600-nucleotide SSU record, QB275330 to the 809- nucleotide ITS record, and QB281636 to the 1,425-nucleotide *rbc*L record of CCAH0201. The homologous analysis windows comprised 38 sequences × 550 columns for SSU-3′, 35 sequences × 711 columns for ITS, and 32 sequences × 1,170 columns for *rbc*L. The *rbc*L window corresponded to positions 61–1230 of its deposited sequence, using one-based numbering.

### 2.5. Alignment and phylogenetic analysis

The three markers were aligned separately with MAFFT v7, and homologous regions were checked manually (Katoh and Standley, 2013). Maximum-likelihood trees were inferred with IQ-TREE 2.4.0; ModelFinder selected models by BIC. ITS used TN+F+I+G4, *rbc*L used TIM+F+G4, and SSU-3′ used K2P. Branch support was assessed with 1,000 SH-aLRT and 1,000 ultrafast bootstrap replicates (Kalyaanamoorthy et al., 2017; Minh et al., 2020; Hoang et al., 2018). Pairwise differences were calculated over columns containing A, C, G, or T in both sequences. Each marker was analyzed in its own matrix.

For a three-marker sensitivity analysis, we retained the 31 taxa present in all three alignments. The homologous SSU-3′ (550 nt), ITS (711 nt), and *rbc*L (1,170 nt) windows were concatenated in that order (2,431 columns) and analyzed with IQ-TREE 2.4.0 under an edge-linked-proportional partition model. ModelFinder selected TNe+I, TIM+F+I+G4, and TIM+F+G4 for the respective partitions by BIC. Branch support was assessed with 1,000 SH-aLRT and 1,000 ultrafast bootstrap replicates; SAG 379-4 was specified as the outgroup. The analysis was repeated with the IQ-TREE -safe option as a numerical sensitivity check using the same matrix and random seed (201).

### 2.6. ITS2 secondary-structure analysis

CCAH0201 ITS2 spans 292 nucleotides within a 340-nucleotide partial 5.8S–ITS2– partial 28S segment with 24 flanking nucleotides on each side; P1 and SAG 2138 were processed in the same way. mfold 3.6 was run with the exact flanks and parameters P=20 and MAX=50 (Zuker, 2003). LocARNA 2.0.0 performed sequence–structure alignment with consistency transformation and produced a maximum expected accuracy (MEA) consensus. The 357-column alignment contained 81 consensus pairs; sequences, CT/dot-bracket structures, alignment and pair map are supplied as supplementary data (Will et al., 2012).

Nucleotide coordinates and base-pair tables were processed with Python 3.13.5 and Biopython 1.86. Sequence, dot-bracket, and CT representations were converted into nucleotide and paired-site tables. Each nucleotide participated in at most one pair, i–j relationships were symmetric, and each pair was counted once with i < j. Pairs were classified as Watson–Crick, G–U/U–G wobble, or noncanonical. P1 mfold structure 6 provided the framework for mapping homologous regions after comparison with the LocARNA/MEA alignment.

Paired positions were mapped through the sequence–structure alignment to identify homologous pairs between CCAH0201 and comparator material. A strict CBC was scored when both nucleotides changed between Watson–Crick states; a hemi-CBC changed one side while retaining pairing potential, including wobble states. Pairs containing gaps or lacking homologous positions were recorded separately. CBC and hemi-CBC classifications used positions with clear structural mapping.

Labels III* and IV* in panel D denote regions mapped by alignment from corresponding elements of P1 mfold structure 6. The five paired-site changes shown in panel D and comparison boxes A–C follow the LocARNA/MEA coordinate table. Each nucleotide, the 5′–3′ direction, and the consensus pairs are retained in the drawing.

## 3. Results

### 3.1. Morphology and cell-size variation of CCAH0201 in the context of related *Stichococcus*-like algae

Of 59 objects screened in the image set, 36 cells met the measurement criteria and represented 22 micrographs. CCAH0201 cells were cylindrical to short-cylindrical, with rounded or blunt ends; some were slightly curved. Solitary cells predominated in the selected fields, while some cells occurred close together. Green intracellular material was unevenly distributed and often offset to one side; bright polar regions and rounded bright bodies appeared in some cells (Fig. 2).

**Figure 2.**
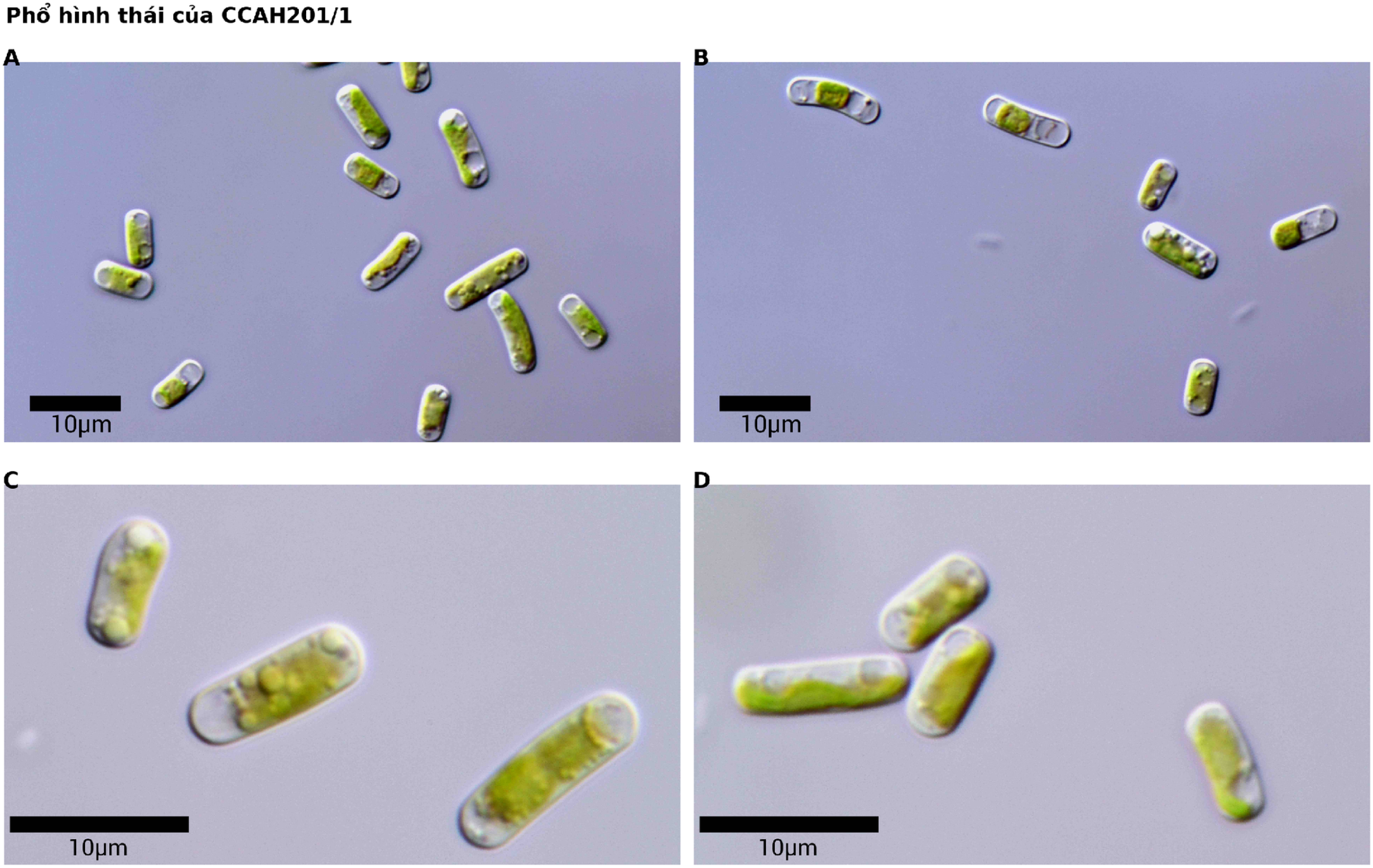
Light micrographs of CCAH0201 with 10-µm scale bars on each source image. Rounded bright bodies are visible in the indicated panels. CCAH201/1 on a source image is a historical alias of CCAH0201/1.

Cell length ranged from 6.68 to 12.74 µm (9.09 ± 1.91 µm; median 8.15 µm), whereas width ranged from 3.49 to 4.79 µm (4.01 ± 0.29 µm; median 4.00 µm). The length-to-width ratio ranged from 1.62 to 3.21, with a mean of 2.26 ± 0.41. Descriptive coefficients of variation were 21.1% for length and 7.2% for width. In blinded within-observer remeasurement of 10 cells, the median absolute difference was 1.39% for length and 7.53% for width. All 10 repeated width measurements exceeded the first measurement, with a mean directional difference of +0.289 µm (+7.16%) and a maximum of +19.06%. Measurement distributions are shown in Fig. S2.

Quantitative comparisons with related material place CCAH0201 within the broad morphological range of *Tetratostichococcus* but reveal no single diagnostic size character. Authenticated *T. jenerensis* CAUP J1302/SAG 2138 has been described as cylindrical with rounded ends, 4.0–11.8 µm long and 2.5–4.2 µm wide. Most of the CCAH0201 range therefore overlaps this material, although its maximum length (12.74 µm) and width (4.79 µm) are greater (Neustupa et al., 2007). *Tetratostichococcus* sp. P1 is smaller under standard shaken culture conditions (3.5–8 × 2–3 µm), although cells can elongate to approximately 30 µm under 1% CO₂, illustrating condition-dependent size plasticity (Sahabudin et al., 2022). *Tetratostichococcus* sp. ACSSI 446 has cylindrical, rounded-ended, occasionally slightly curved cells 4–19 µm long and 2.5–3.9 µm wide, with a mean of 7.8 × 3.2 µm. Its length range broadly overlaps that of CCAH0201, whereas CCAH0201 has a somewhat greater mean and maximum width (Tukhtaboeva et al., 2024).

Cell-size ranges also overlap with *Tritostichococcus* corticulus, whose cells are 6.3– 10.1 µm long and 3.1–4.1 µm wide, with a length-to-width ratio of 1.7–2.9 (Pröschold and Darienko, 2020). Cellular organization provides additional comparative characters: *T. jenerensis* has a parietal chloroplast with a starch-enveloped pyrenoid and may form short filaments, whereas ACSSI 446 sometimes forms two-cell filaments and has a pyrenoid. CCAH0201 dimensions and shape describe a Stichococcus-like morphotype; molecular markers further resolve its taxonomic position.

### 3.2. Conserved SSU-3′ and discriminatory ITS and *rbcL* signals

The analyzed CCAH0201 material comprised a 550-nucleotide SSU-3′ segment, an 809-nucleotide ITS consensus, and a 1,425-nucleotide *rbcL* fragment. The ITS consensus contained the 292-nucleotide ITS2 region modeled below. The *rbcL* fragment comprised 475 codons and had no internal stop codon in the analyzed reading frame; the homologous tree window retained 1,170 columns.

CCAH0201, P1 and SAG 2138 shared the same nucleotide states across the 550 compared SSU-3′ positions. The ML tree of 38 sequences × 550 columns therefore provides a conserved-marker context for the ITS and *rbc*L comparisons (Fig. S1).

ITS and *rbcL* showed greater divergence than SSU-3′, and CCAH0201 was closer to P1 than to SAG 2138 within the homologous range of each matrix (Table 2). Nine uncovered *rbc*L columns in SAG 2138 were excluded from the pairwise denominator. In the SSU-3′ ML tree, 14 very short internal branches (<0.0018 substitutions/site) correspond to the low variation in this region (Fig. S1).

**Table 1.** Quantitative morphometry of 36 CCAH0201 cells from 22 micrographs. Here, n is the number of observed cells, not the number of independent biological replicates.

| **Variable** | **Observed range** | **Mean ± SD** | **Median; Q1–Q3** |
|---|---|---|---|
| Cell length, µm | 6.68–12.74 | 9.09 ± 1.91 | 8.15; 7.54–10.98 |
| Cell width, µm | 3.49–4.79 | 4.01 ± 0.29 | 4.00; 3.83–4.19 |
| Length-to-width ratio | 1.62–3.21 | 2.26 ± 0.41 | 2.15; 1.94–2.49 |

**Table 2.** Single-locus differences between CCAH0201 and P1 or SAG 2138.

| **Comparison region** | **Comparator** | **Substitutions/comparable ACGT sites** | **Substitution proportion, %** |
|---|---|---|---|
| SSU-3′ | P1 | 0/550 | 0.000 |
| SSU-3′ | SAG 2138 | 0/550 | 0.000 |
| ITS | P1 | 22/711 | 3.094 |
| ITS | SAG 2138 | 60/711 | 8.439 |
| *rbcL* | P1 | 4/1,170 | 0.342 |
| *rbcL* | SAG 2138 | 28/1,161 | 2.412 |

The ITS ML tree placed CCAH0201 and P1 in the same group, with SH-aLRT/UFBoot support of 87/94 (Fig. 3).

**Figure 3.**
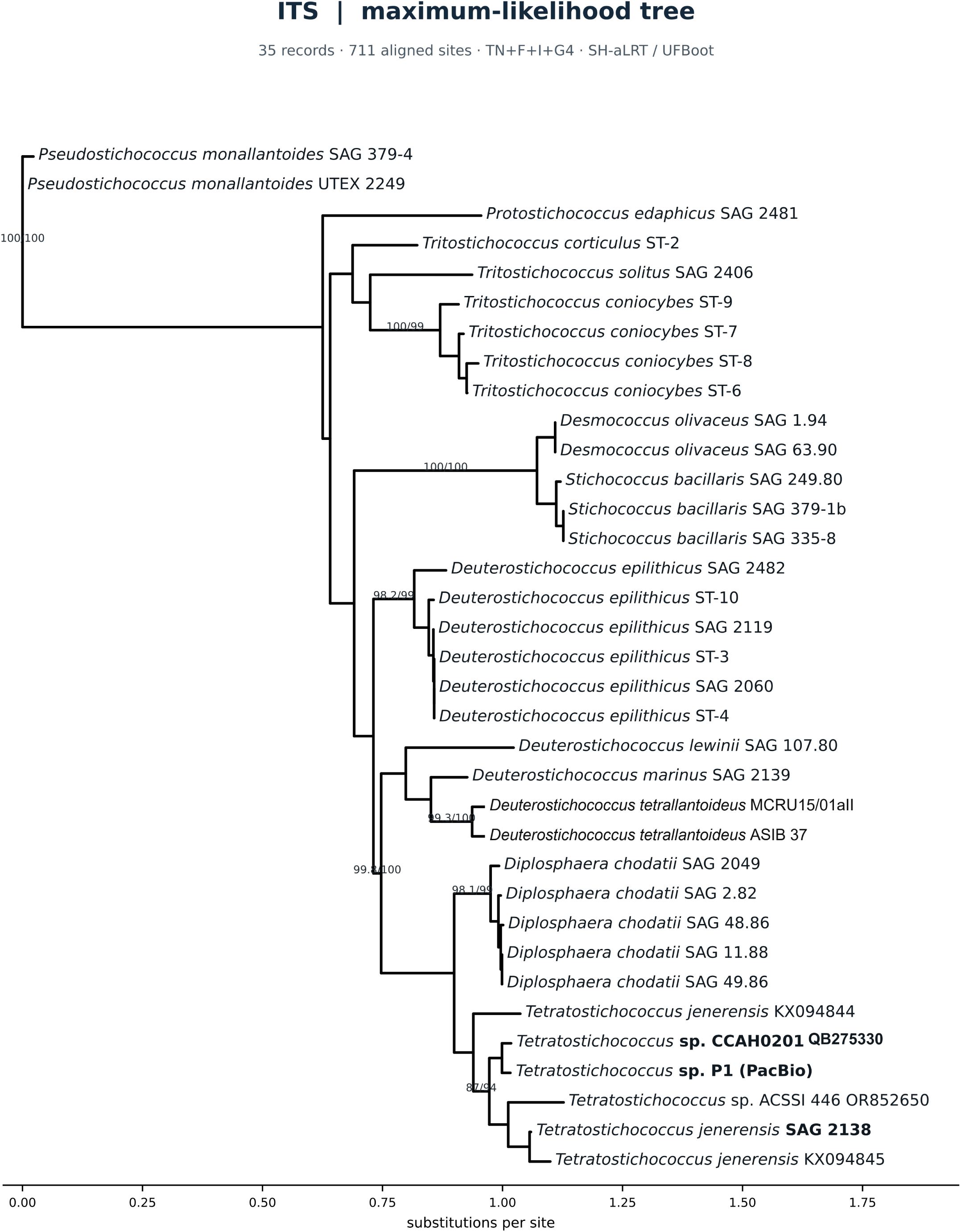
Maximum-likelihood ITS tree inferred from 35 records × 711 columns under TN+F+I+G4 with IQ-TREE 2.4.0. The CCAH0201 tip bears GenBank accession QB275330 for the 809-nucleotide ITS record; the tree uses a 711-column homologous window. Branch values indicate SH-aLRT/UFBoot support (%) from 1,000 replicates each. The CCAH0201–P1 group has support of 87/94. The scale bar denotes substitutions per site. The tree is oriented for display on the *Pseudostichococcus monallantoides* edge (SAG 379-4 and UTEX 2249).

The *rbcL* ML tree also placed CCAH0201 close to P1, with SH-aLRT/UFBoot support of 96.9/99 (Fig. 4).

**Figure 4.**
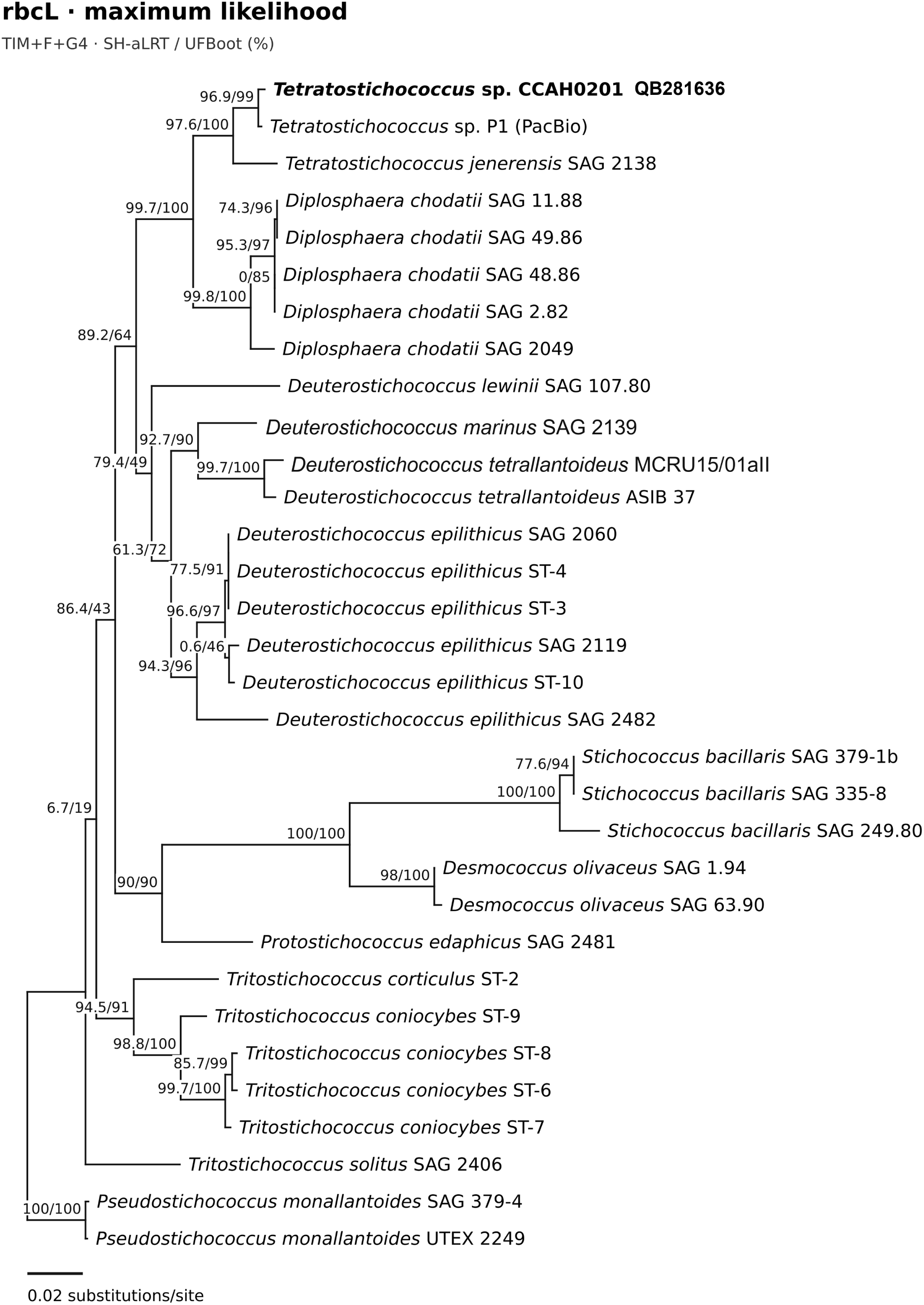
Maximum-likelihood *rbc*L tree inferred from 32 records × 1,170 columns under TIM+F+G4 with IQ-TREE 2.4.0. The CCAH0201 tip bears GenBank accession QB281636 for the 1,425-nucleotide *rbc*L record; the tree uses a 1,170-column homologous window. Branch values indicate SH- aLRT/UFBoot support (%) from 1,000 replicates each. The CCAH0201–P1 group has support of 96.9/99. The scale bar denotes substitutions per site. The tree is oriented for display on the *Pseudostichococcus monallantoides* edge (SAG 379-4 and UTEX 2249).

In the 31-taxon concatenated analysis, CCAH0201 and P1 formed a group with SH- aLRT/UFBoot support of 98.7/100. That pair grouped with *T. jenerensis* SAG 2138 with support of 99.5/100 (Fig. S3). The standard and -safe runs yielded the same topology and support labels. Both produced numerical underflow warnings during ModelFinder, before the final tree search (Fig. S3).

### 3.3. ITS2 structural variation among CCAH0201, P1, and SAG 2138

Figure 5 presents consensus ITS2 models for CCAH0201 (292 nucleotides), P1 (301 nucleotides) and SAG 2138 (314 nucleotides), each showing 81 pairs from the LocARNA/MEA alignment. CCAH0201–P1 had one strict CBC (UA→CG), one two-sided change originating from a wobble pair (UG→GC) and three hemi-CBCs. CCAH0201–SAG 2138 had four strict CBCs and five hemi-CBCs. ITS2 ranges for each material are given in Table 3.

**Figure 5.**
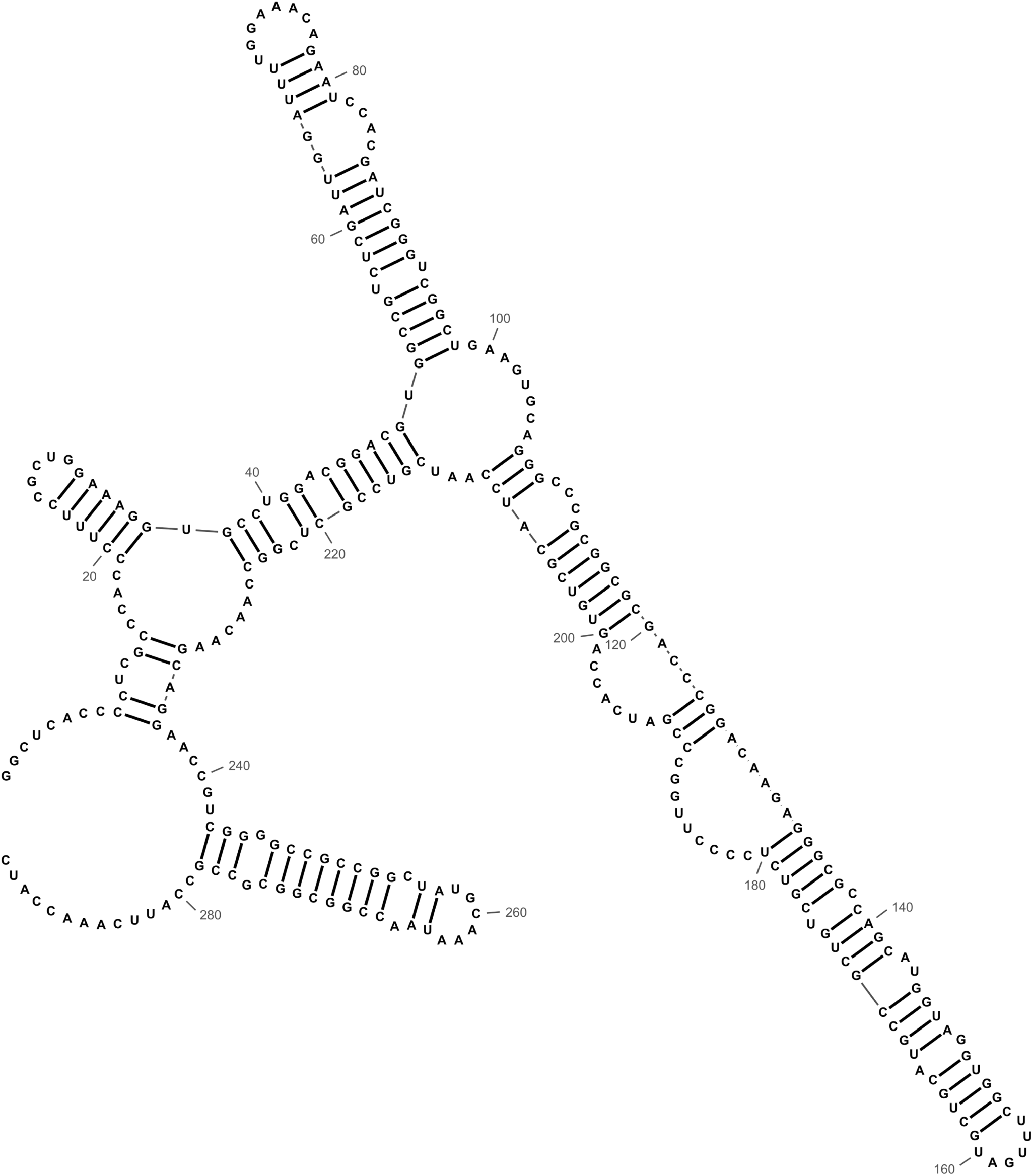
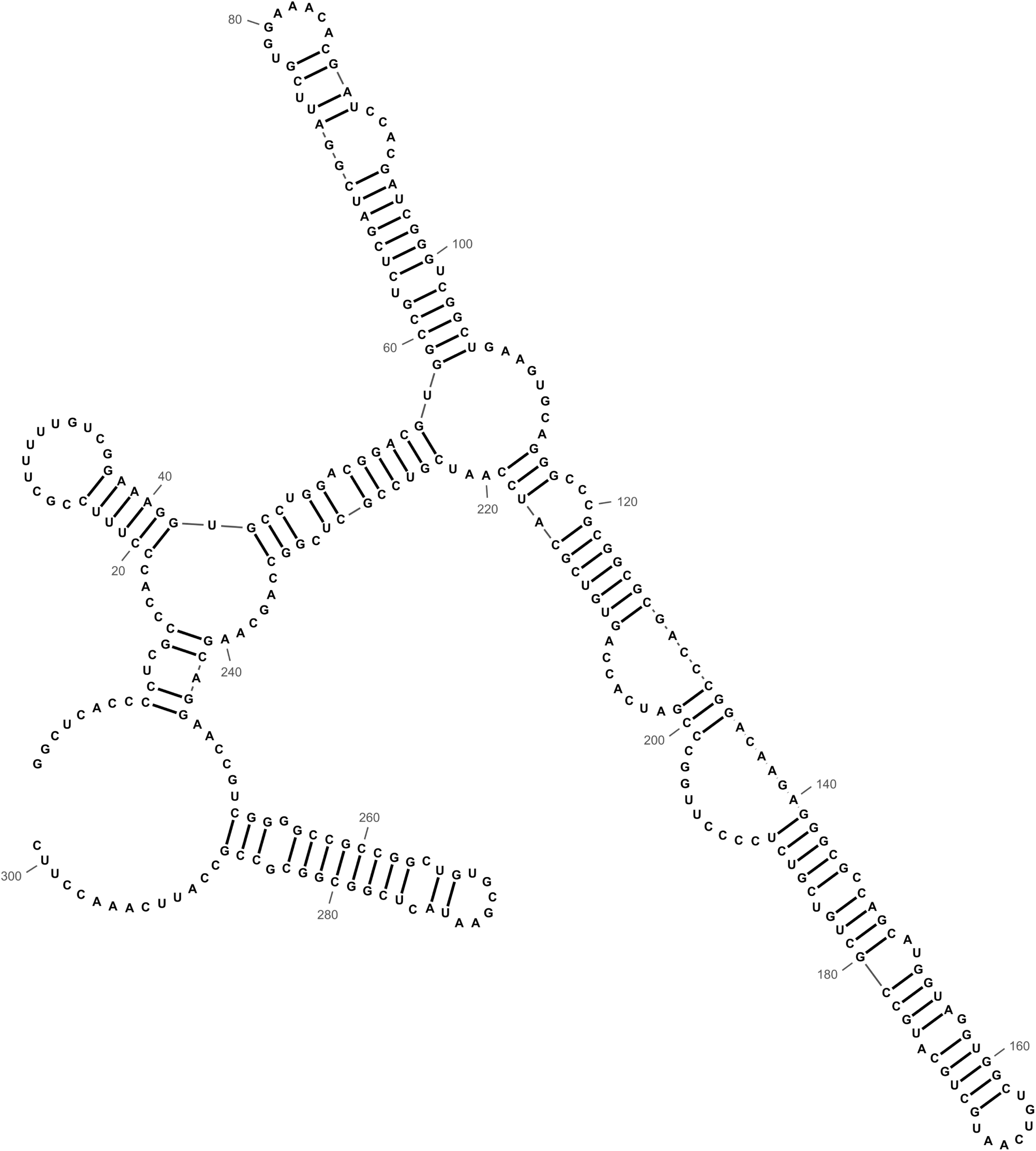
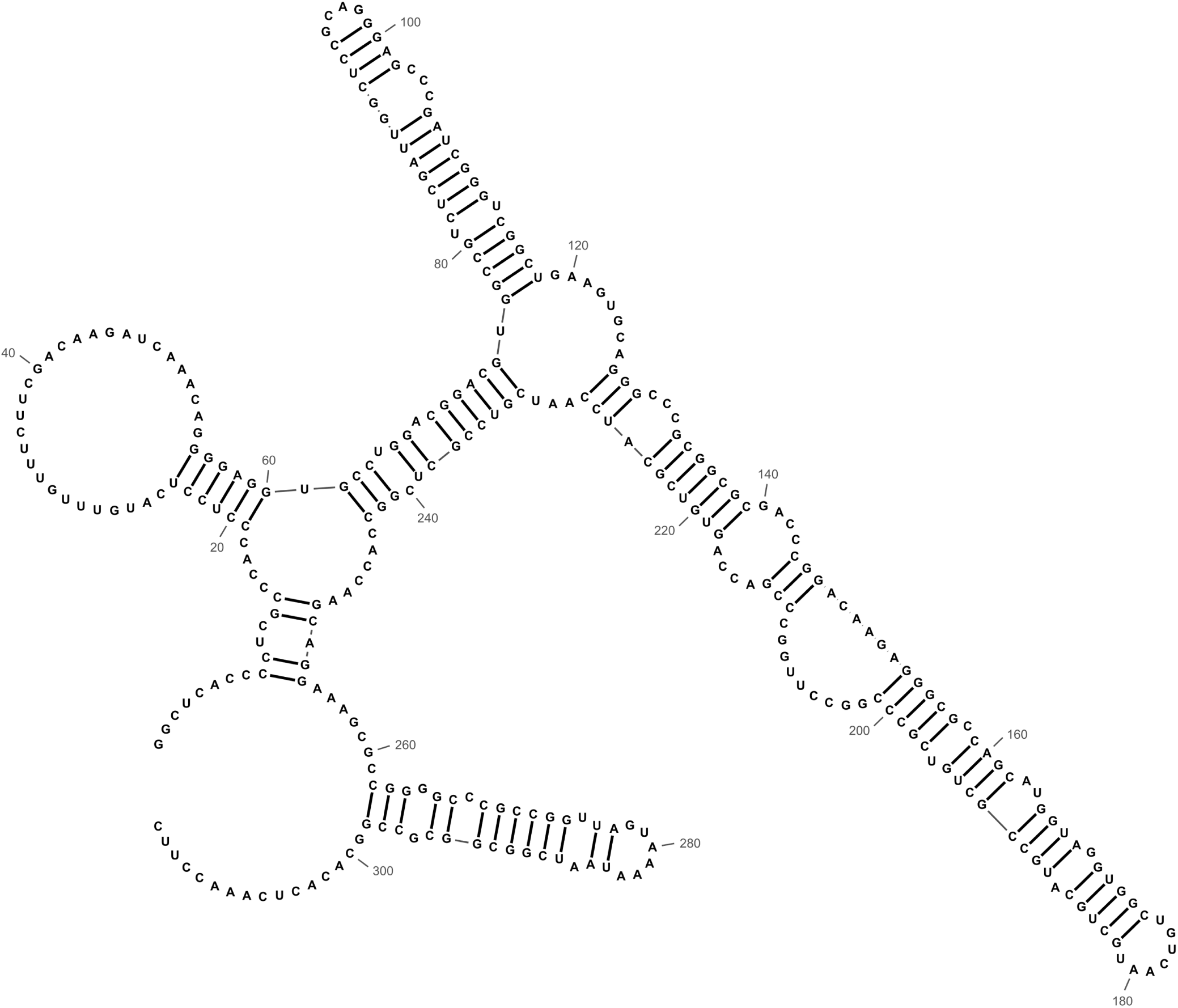
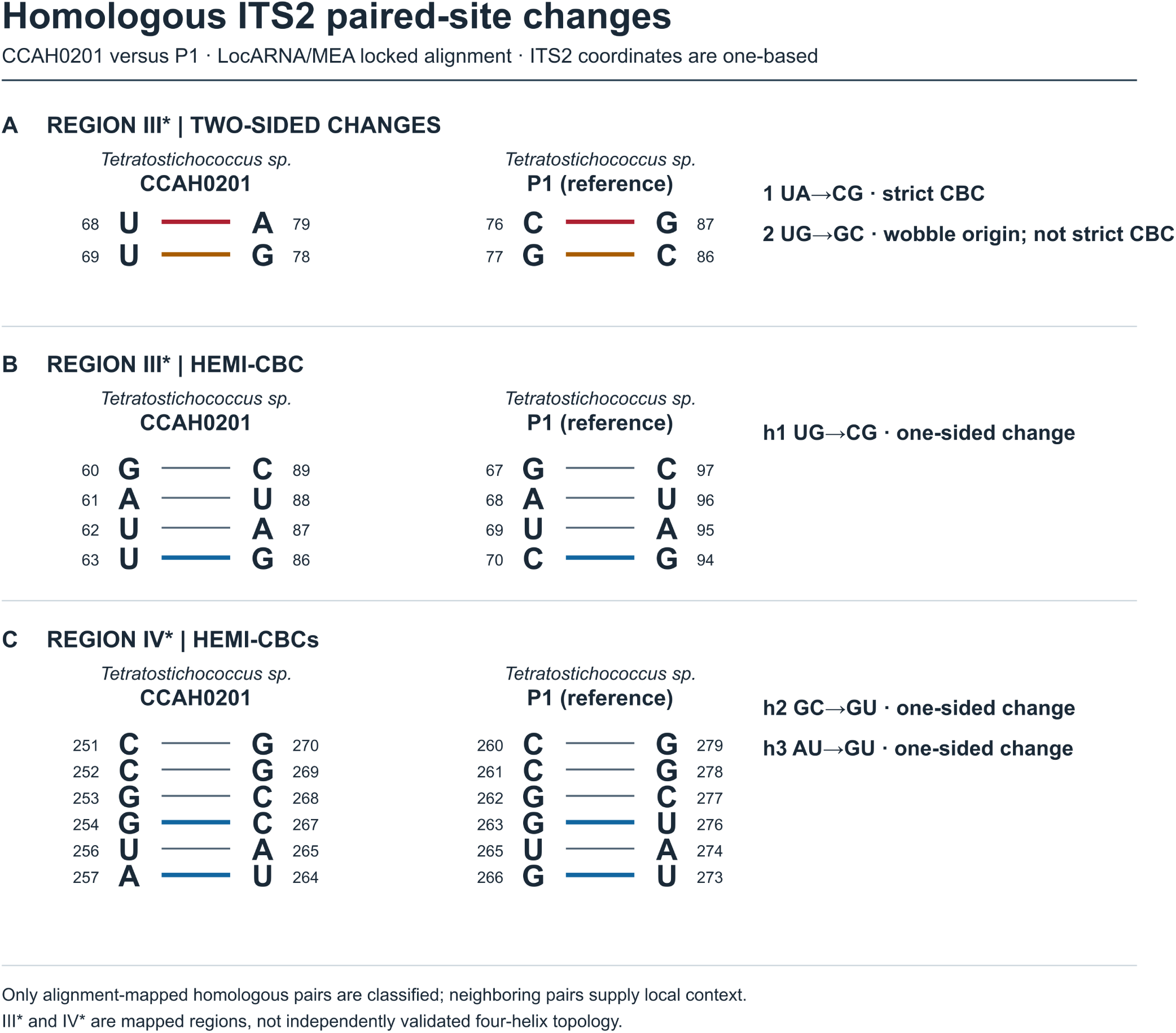
Vector renderings of consensus ITS2 secondary-structure models, laid out in RNAcanvas (Johnson and Simon, 2023). Each panel displays 81 pairs from the LocARNA/MEA sequence–structure alignment. P1 mfold structure 6 guided mapping of regions III*–IV* in panel D. (A) *Tetratostichococcus* sp. CCAH0201, 292 nucleotides. (B) *Tetratostichococcus* sp. P1, 301 nucleotides. (C) *T. jenerensis* SAG 2138, 314 nucleotides. (D) Five homologous paired-site changes between CCAH0201 and P1: one strict CBC UA→CG, one two-sided wobble-origin change UG→GC and three hemi-CBCs. Coordinates are one-based in each sequence; asterisks mark alignment-mapped regions.

**Table 3.** ITS2 boundaries within the sequence segments used for structural modelling.

| **Material** | **Full-segment length (nt)** | **ITS2 start** | **ITS2 end** | **ITS2 length (nt)** |
|---|---|---|---|---|
| *Tetratostichococcus* sp. CCAH0201 | 340 | 25 | 316 | 292 |
| *Tetratostichococcus* sp. P1 | 349 | 25 | 325 | 301 |
| *T. jenerensis* SAG 2138 | 362 | 25 | 338 | 314 |
| *T. jenerensis* KS126SM6L | 381 | 25 | 357 | 333 |

Coordinates are one-based and inclusive. Full-segment length includes ITS2 and flanking nucleotides. Table 4 compares the 81 consensus pairs within ITS2; full- segment models have different scopes and denominators.

**Table 4.**
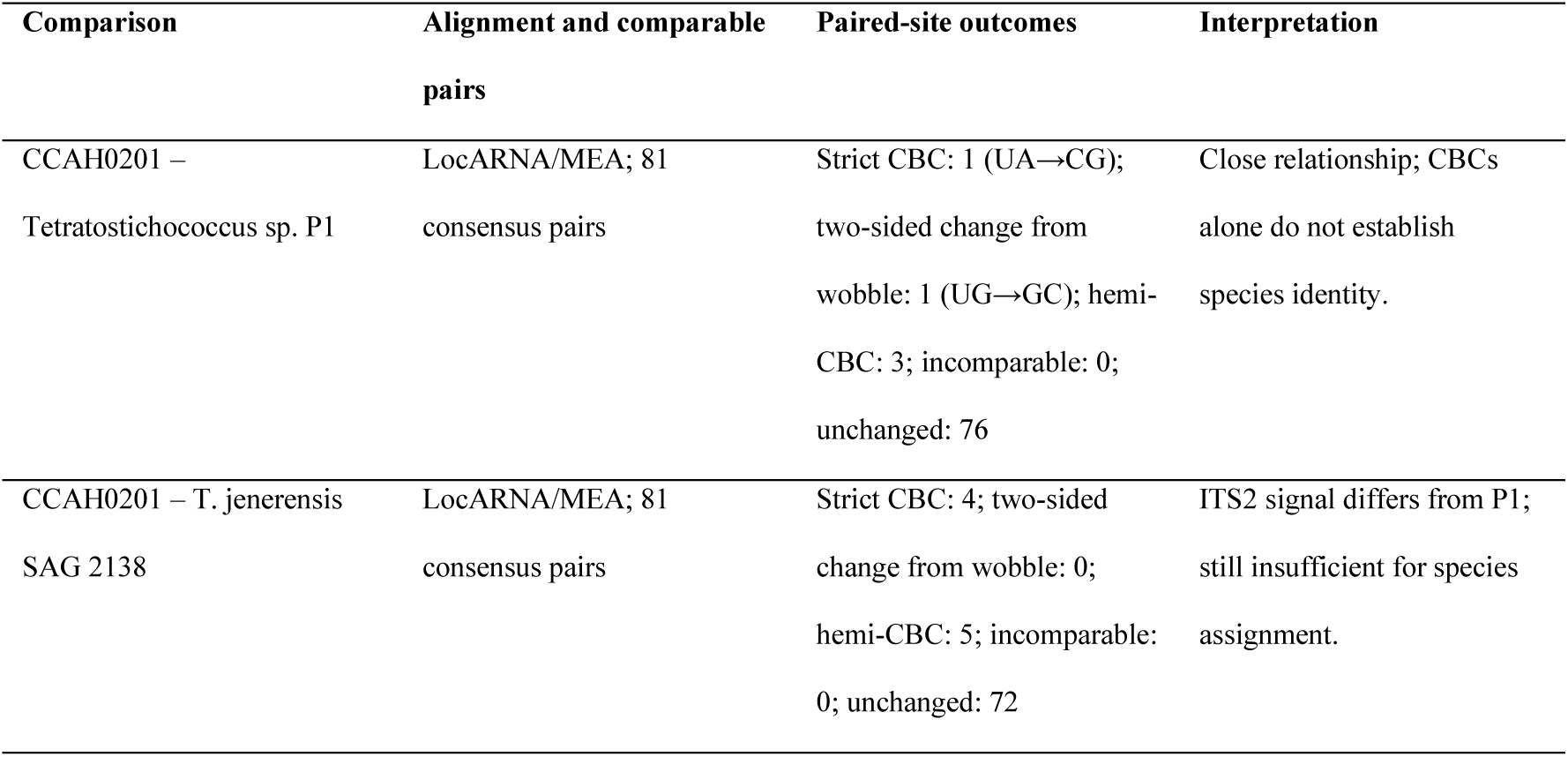
Paired-site changes among 81 consensus ITS2 pairs in the LocARNA/MEA alignment.

### 3.4. Integrative identification

CCAH0201 morphology is consistent with *Stichococcus*-like algae. SSU-3′ places the strain in the *Tetratostichococcus* context, while ITS and *rbcL* consistently show a close relationship to P1 in the ML analyses. Differences from SAG 2138 occur in ITS and *rbcL* sequences and in ITS2 paired sites. Together, these comparisons support the designation *Tetratostichococcus* sp. CCAH0201 and identify a specific candidate for further lineage-delimitation research (Table 5).

**Table 5.**
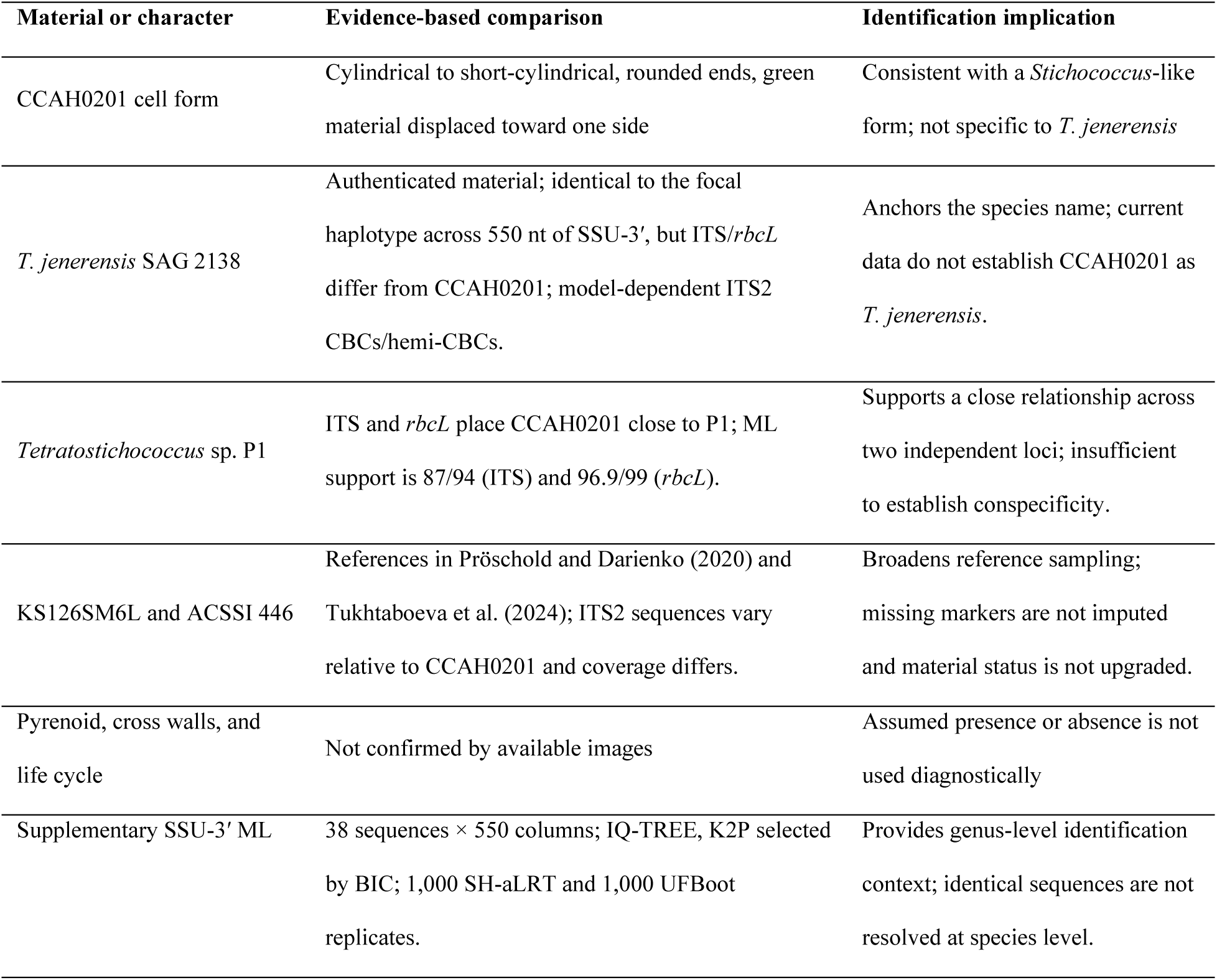
Diagnostic comparisons and identification limits for CCAH0201.

| Material or character | Evidence-based comparison | Identification implication |
|---|---|---|
| CCAH0201 cell form | Cylindrical to short-cylindrical, rounded ends, green material displaced toward one side | Consistent with a *Stichococcus*-like form; not specific to *T. jenerensis* |
| *T. jenerensis* SAG 2138 | Authenticated material; identical to the focal haplotype across 550 nt of SSU-3′, but ITS/*rbcL* differ from CCAH0201; model-dependent ITS2 CBCs/hemi-CBCs. | Anchors the species name; current data do not establish CCAH0201 as *T. jenerensis*. |
| *Tetratostichococcus* sp. P1 | ITS and *rbcL* place CCAH0201 close to P1; ML support is 87/94 (ITS) and 96.9/99 (*rbcL*). | Supports a close relationship across two independent loci; insufficient to establish conspecificity. |
| KS126SM6L and ACSSI 446 | References in Pröschold and Darienko (2020) and Tukhtaboeva et al. (2024); ITS2 sequences vary relative to CCAH0201 and coverage differs. | Broadens reference sampling; missing markers are not imputed and material status is not upgraded. |
| Pyrenoid, cross walls, and life cycle | Not confirmed by available images | Assumed presence or absence is not used diagnostically |
| Supplementary SSU-3′ ML | 38 sequences × 550 columns; IQ-TREE, K2P selected by BIC; 1,000 SH-aLRT and 1,000 UFBoot replicates. | Provides genus-level identification context; identical sequences are not resolved at species level. |

## 4. Discussion

### 4.1. A lineage close to P1 identified from *Rhizophora* leaves in Can Gio

The 550-nucleotide SSU-3′ window is identical in CCAH0201, P1 and *T. jenerensis* SAG 2138, so it offers little resolution within this group. ITS and plastid *rbc*L instead place CCAH0201 with P1. The observed cell form supports a *Stichococcus*-like morphotype, while the two informative loci and homologous ITS2 pairs support identification as *Tetratostichococcus* sp.

Reference choice is central to this identification. The revision of *Stichococcus*-like algae by Pröschold and Darienko (2020) established the genus framework, and the independently characterized P1 culture supplies a closer comparator than SAG 2138 (Sahabudin et al., 2022; Maeda et al., 2024). Comparing CCAH0201 with both P1 and the type-species material separates genus placement from the unresolved lineage boundary between CCAH0201 and P1.

### 4.2. Differentiation in ITS, *rbcL*, and ITS2 paired sites

P1 is the closest sampled comparator in both informative markers. CCAH0201 differs from P1 at 22/711 ITS positions and 4/1,170 *rbc*L positions, versus 60/711 and 28/1,161 positions relative to SAG 2138 (Table 2). The CCAH0201–P1 branch receives SH-aLRT/UFBoot support of 87/94 in ITS and 96.9/99 in *rbc*L. Agreement between nuclear and plastid markers strengthens the inferred relationship; these distances alone do not delimit a species.

The partitioned analysis of 31 taxa shared across all three matrices also groups CCAH0201 with P1 (98.7/100) and places that pair with SAG 2138 (99.5/100). Because the SSU-3′ segment is identical across the three focal strains and model selection retained numerical warnings, this tree is treated as a sensitivity check alongside the separate marker trees.

Mapping ITS2 onto 81 homologous consensus pairs yields one strict CBC (UA→CG), one two-sided change from a G–U wobble (UG→GC) and three hemi-CBCs between CCAH0201 and P1. Comparison with SAG 2138 yields four strict CBCs and five hemi- CBCs (Fig. 5; Table 4). The paired-site ranking agrees with the ITS and *rbc*L comparisons.

The starting pair state determines the event class: UG→GC changes both bases but is not counted as a strict CBC because it begins with a wobble pair. Homologous ITS2 sites may show homoplasy or changes on internal branches (Caisová et al., 2011, 2013). The paired-site events therefore inform interpretation of the gene trees rather than serving as a numerical species threshold.

### 4.3. Leaf-derived material in the context of terrestrial diversity

The measured CCAH0201 range of 6.68–12.74 × 3.49–4.79 µm overlaps with *T. jenerensis* SAG 2138/CAUP J1302, ACSSI 446 and *Tritostichococcus corticulus*. Its cylindrical to short-cylindrical morphotype supplies a useful culture description, while ITS, *rbc*L and ITS2 provide the discriminating comparisons (Fig. 2; Table 1; Fig. S2). A survey of tropical rainforest leaves found that phyllosphere algal composition varied with forest type and host tree (Liu et al., 2023). Community surveys and cultured isolates address different scales: CCAH0201 links a living culture from a recorded leaf collection to measured cells and accessioned markers. It can therefore serve as a taxonomic reference when related sequences are recovered from other substrates. CCAH0201 came from a *Rhizophora* leaf, whereas its closest sampled comparator, P1, was isolated from tropical peat soil in Malaysia (Sahabudin et al., 2022). Closely related cultures have thus been recovered from contrasting terrestrial substrates. P1 grew across pH 3–8 in laboratory culture, with the highest measured growth rate at pH 3 (Sahabudin et al., 2022); these physiological measurements characterize P1. The present evidence establishes the placement of CCAH0201 without assigning it a substrate preference or pH response.

The leaf–peat comparison raises a question about how diversity within *Tetratostichococcus* is distributed among terrestrial microhabitats. Tested *Stichococcus*- like genera differed in salt response, and their responses to salt and drying were not coupled (Van et al., 2021). The CCAH0201 culture and marker records make this lineage recognizable in later habitat surveys without relying on cell shape alone.

### 4.4. From marker evidence to tests with comparative genomics

A separately reported whole-genome sequencing (WGS) study of quality-checked CCAH0201 material is planned. The present study assesses its relationship to P1 with SSU, ITS, *rbc*L, ITS2 structure and morphology.

P1 already has a chromosome-scale reference genome (Maeda et al., 2024). Future CCAH0201 sequence data could test whether the relationship inferred here extends to single-copy nuclear genes and organellar genomes. Comparisons that include name- linked *T. jenerensis* SAG 2138 and other close strains would place variation between CCAH0201 and P1 in a broader genus context. Genomic similarity by itself would not define a species boundary or establish an ecological trait.

### 4.5. Study limitations

This study includes one CCAH0201 strain and the currently available reference set; affinity to P1 does not establish a species boundary. The 550-nucleotide SSU-3′ window has little variation in the focal group. ITS2 structures are predictions, and the ITS2 boundaries were defined from the modelled sequence rather than independently annotated. The 81-pair consensus projection does not retain an intraregional stem in region I, so complete four-helix topology requires further testing. The concatenated analysis is limited to 31 common taxa and retained numerical warnings during model selection.

The micrographs capture one culture time point. Cells close together in still images do not show whether filaments develop, and culture conditions at imaging were insufficient for analysis of phenotype across environments. Repeat measurements showed a mean directional width difference of +7.16%. Bright rounded bodies require targeted observation before assignment as pyrenoids or reproductive structures. Salt tolerance and leaf specialization likewise require direct assessment.

## 5. Conclusions

CCAH0201 from *Rhizophora* leaves in the Can Gio Mangrove Forest, Vietnam, is identified as *Tetratostichococcus* sp. in a lineage close to P1. Nuclear ITS and plastid *rbc*L support this relationship, while the SSU-3′ window does not resolve the focal group. ITS2 adds structural evidence in comparisons with P1 and SAG 2138. The culture, quantitative morphological record, and three marker regions provide a reference for surveying diversity within the genus. Additional strains, standardized culture conditions, and comparative genomic data can test lineage boundaries and related biological differences.

## Supplementary material

The SSU-3′ tree (Fig. S1) and CCAH0201 morphometry distributions (Fig. S2) are supplied as supplementary figures. The accompanying data include the marker alignments and trees, pairwise comparisons, ITS2 structures and coordinate tables, and morphometry measurements. The 31-taxon partitioned tree (Fig. S3), matrix, partition file, and IQ-TREE output are also supplied.

## Data and code availability

GenBank assigned accession numbers QB266712 (SSU, 600 nucleotides), QB275330 (ITS, 809 nucleotides), and QB281636 (*rbc*L, 1,425 nucleotides) to the CCAH0201 sequences. The phylogenetic trees use homologous windows of 550, 711, and 1,170 nucleotides, respectively. Alignments, tree files, ITS2 pair maps and coordinates, and morphometry data are provided as supplementary data. The supplementary data also contain the 31-taxon three-marker matrix, partition definitions, Newick tree, model report, and analysis log. The versioned data and analysis code are openly available at Zenodo: https://doi.org/10.5281/zenodo.23969895.

## Acknowledgements

This study received no dedicated funding. The authors declare no competing interests. OpenAI Codex assisted with English-language editing and preparation of analysis and figure-rendering scripts. The concatenated analysis matrix, scripts, and output files are supplied as supplementary material.

## Supplementary figure legends

**Figure S1.**
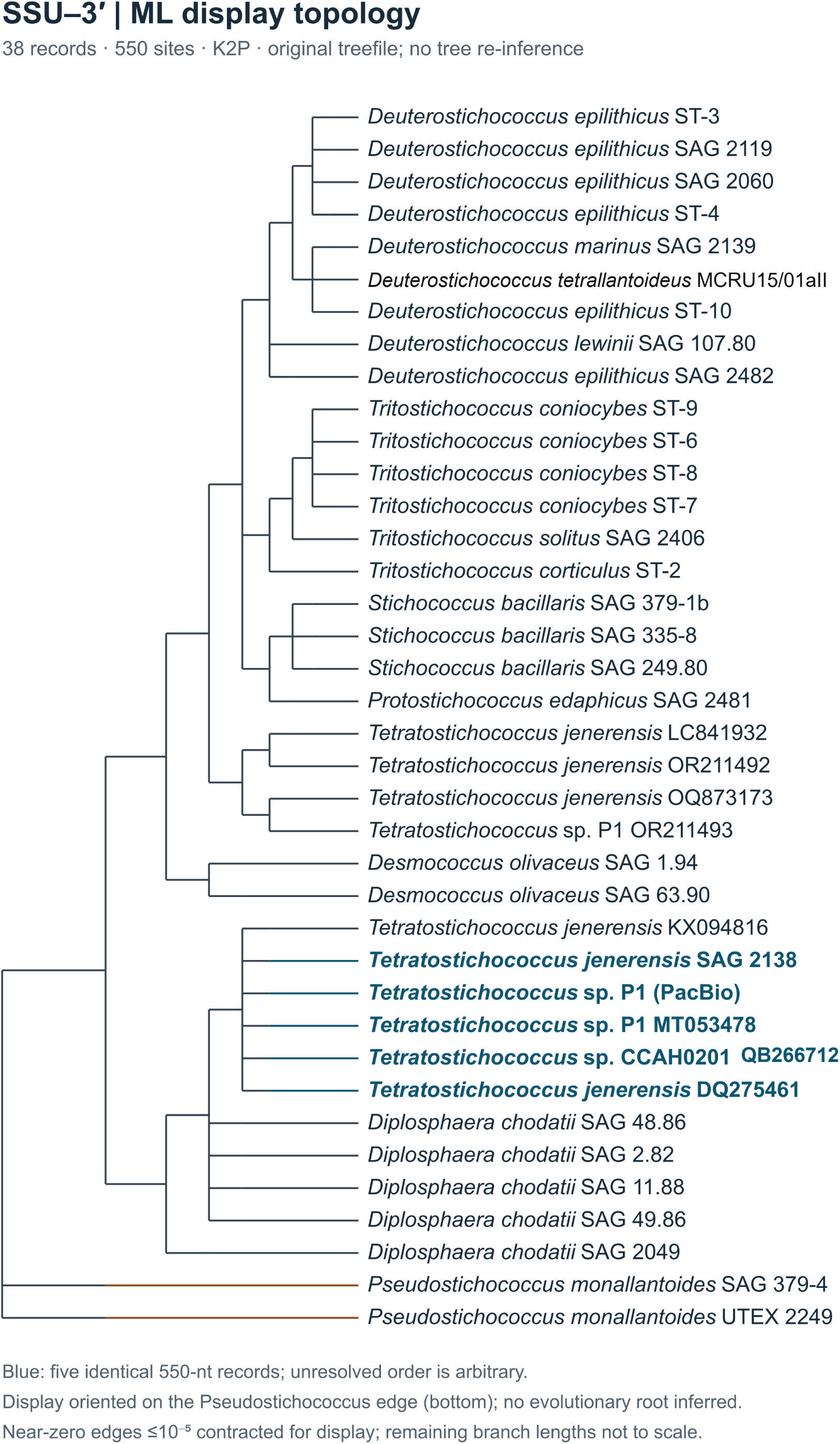
SSU-3′ maximum-likelihood tree inferred from 38 records × 550 columns under K2P. The CCAH0201 tip bears GenBank accession QB266712 for the 600- nucleotide SSU record; the tree uses a 550-column homologous window. Very short edges (≤10⁻⁵ substitutions/site) were contracted for display. Five blue-highlighted records have identical sequences across the analysis window. The tree is oriented for display on the *Pseudostichococcus monallantoides* edge (SAG 379-4 and UTEX 2249); remaining edge lengths are not to scale.

**Figure S2.**
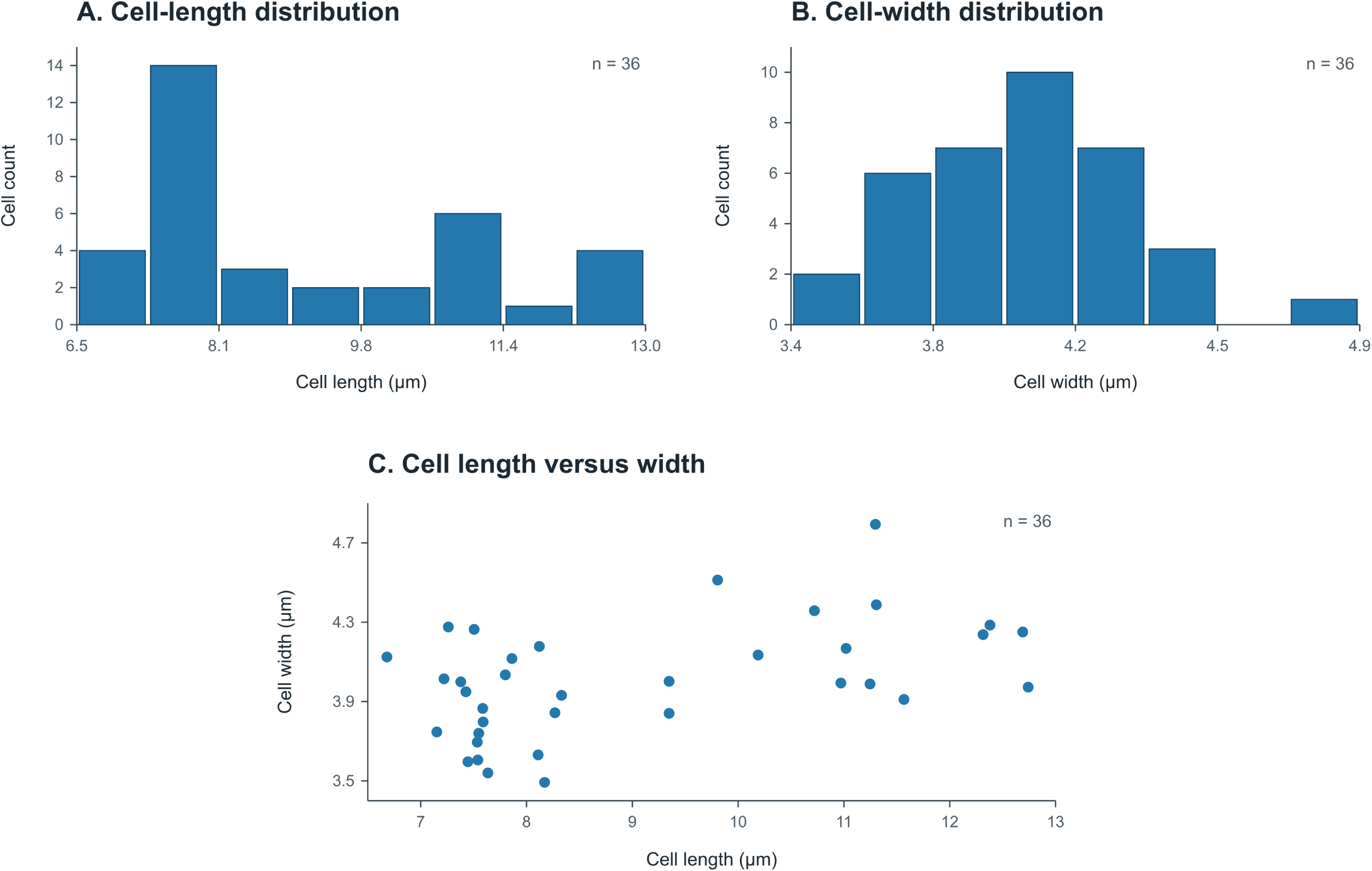
Morphometry distributions for 36 individual CCAH0201 cells. (A) Cell- length distribution; (B) cell-width distribution; (C) relationship between length and width for each cell. Values were converted from the 10-µm scale bars of the corresponding image group and use the same measurements summarized in Table 1.

**Figure S3.**
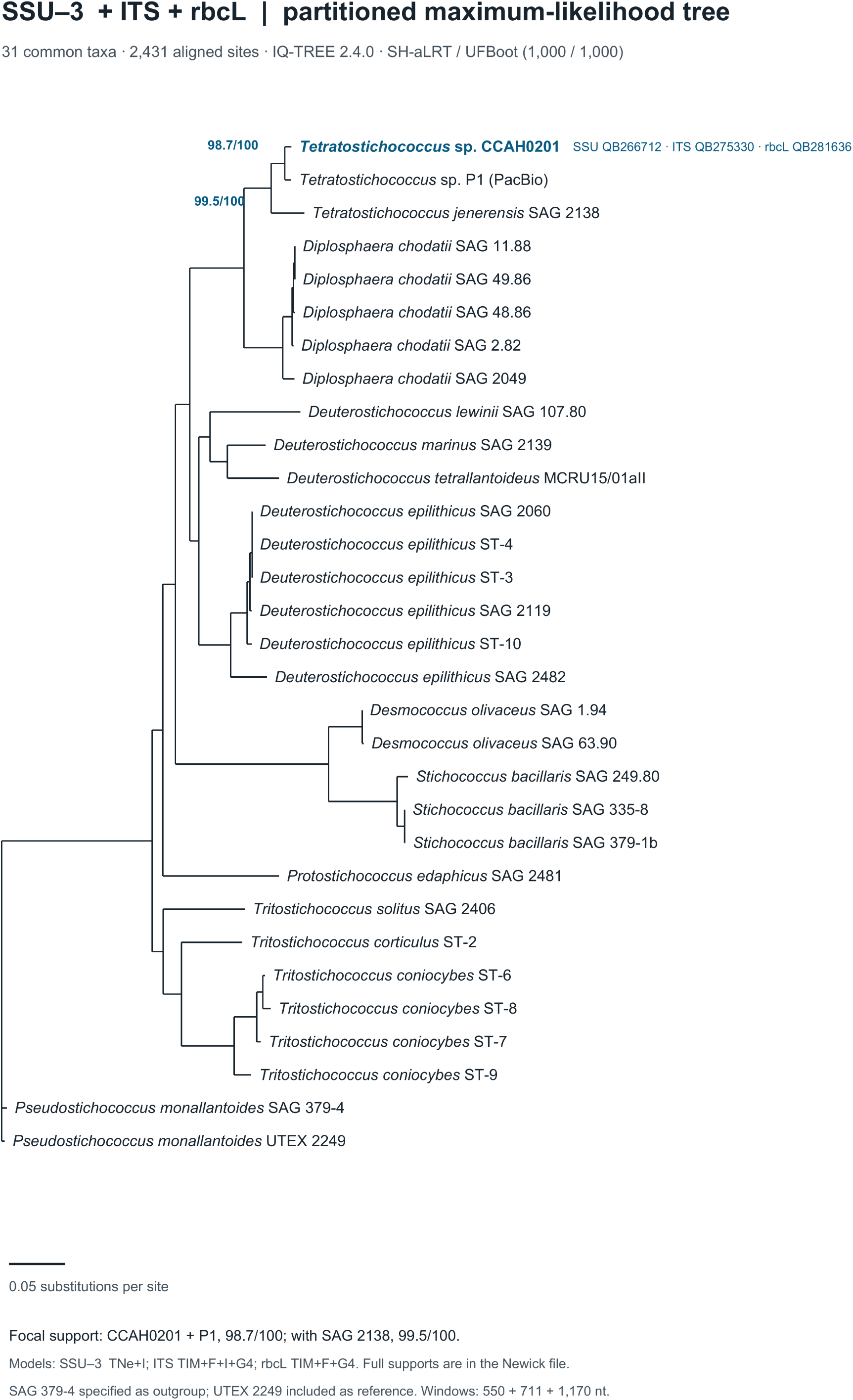
Partitioned maximum-likelihood tree of 31 taxa inferred from SSU-3′ (550 nt), ITS (711 nt), and *rbc*L (1,170 nt) windows (2,431 aligned columns). IQ-TREE 2.4.0 selected TNe+I, TIM+F+I+G4, and TIM+F+G4 by BIC for the three partitions. Only support for the CCAH0201–P1 pair (98.7/100) and its group with *T. jenerensis* SAG 2138 (99.5/100) is printed on the figure; values are SH-aLRT/UFBoot percentages from 1,000 replicates each. Other support values and branch lengths are in the supplied Newick tree. SAG 379-4 was specified as the outgroup, and the scale bar denotes 0.05 substitutions per site. The CCAH0201 tip shows GenBank accessions QB266712 (SSU), QB275330 (ITS), and QB281636 (*rbc*L). The MH670366.1/MH672651.1 tip is shown as *D. tetrallantoideus* (voucher MCRU15/01aII; deposited as *S. antarcticus*; Pröschold, Darienko and Guiry, 2020).

